# Promoter replacement by genome editing creates gain-of-function traits in Arabidopsis

**DOI:** 10.1101/2025.03.22.643860

**Authors:** Takashi Nobusawa, Michiharu Nakano, Yumi Nagashima, Makoto Kusaba

**Affiliations:** Graduate School of Integrated Sciences for Life, Hiroshima University, 1-4-3, Kagamiyama, Higashi-Hiroshima 739-8526, Japan; Faculty of Agriculture and Marie Science, Kochi University

**Keywords:** SDN-1, non-GMO, inversion, translocation, chromosome

## Abstract

We introduced two targeted DNA double-strand breaks on the same Arabidopsis chromosome using CRISPR-Cas9 and replaced the *FLOWERING LOCUS T* (*FT*) promoter with that of a histone variant gene through chromosomal inversion. The resulting lines misexpressed *FT* and flowered early, like *FT*-overexpressing transgenic plants. This system can be used to create gain-of-function mutations that modify target gene expression as desired without incorporating foreign DNA sequences.

---

Modifying target genes through genome editing technology has promising agricultural applications (Wang et al., 2023). Typical genome editing employs site-directed nucleases (SDNs), such as clustered regularly interspaced palindromic repeats (CRISPR)-associated protein 9 (Cas9), to generate DNA double-strand breaks (DSBs) at specific sites in the genome, leading to DNA mutations caused by errors during repair. The mutations generated by genome editing using SDNs can be classified into three categories (Matres et al., 2021). SDN-1 mutations are typically short deletions or insertions, arising through errors during DNA repair via non-homologous end-joining. SDN-2 and -3 mutations result from homology-directed repair, whereby foreign DNA sequences are introduced into the genome DNA. Even though genome-edited plants carrying SDN-1 type mutations were produced through transgenesis, they are not heavily regulated and are socially acceptable in many countries as long as their transgenes have been segregated out. Therefore, SDN-1 genome editing is useful for crop improvement, and many SDN-1 genome-edited strains have been developed. However, most SDN-1 genome-edited strains developed so far harbor loss-of-function mutations.

Two DSBs on the same chromosome may cause a deletion or an inversion. Such inversions have been reported to be achieved in plants using CRISPR-Cas9 (Schmidt et al., 2019). Because such modifications do not involve foreign DNA sequences, they are considered to be SDN-1 type mutations. Most induced inversions cause loss-of-function mutations. However, an inversion that creates a gene fusion involving the coding region of the target gene and the promoter of another gene may affect the expression of the target gene, leading to phenotypic changes.

To test this possibility, we aimed to exchange the promoters of two genes on the same chromosome via targeted inversion in the model plant Arabidopsis (*Arabidopsis thaliana*), using CRISPR-Cas9 to introduce DSBs near the transcription start sites. We chose *FLOWERING LOCUS T* (*FT*) on chromosome 1 as the target gene. *FT* encodes florigen and leads to very early flowering when overexpressed (Kardailsky et al., 1999). As a promoter donor, we used *HTA3*, encoding a histone H2A variant. *HTA3* is expressed in various young tissues, and its loss-of-function mutants exhibit normal growth (Yi et al., 2006). *HTA3* is also located on chromosome 1, about 3.6 Mb upstream of *FT*, with the two genes in divergent orientations. Thus, an inversion of the 3.6-Mb fragment between the two genes that contains their promoters would result in a promoter exchange, placing *FT* under the control of the *HTA3* promoter. We therefore designed two single guide RNAs (sgRNAs) to introduce DSBs near the transcription start sites of *FT* and *HTA3* (Fig. 1a, b).

**Figure 1.**
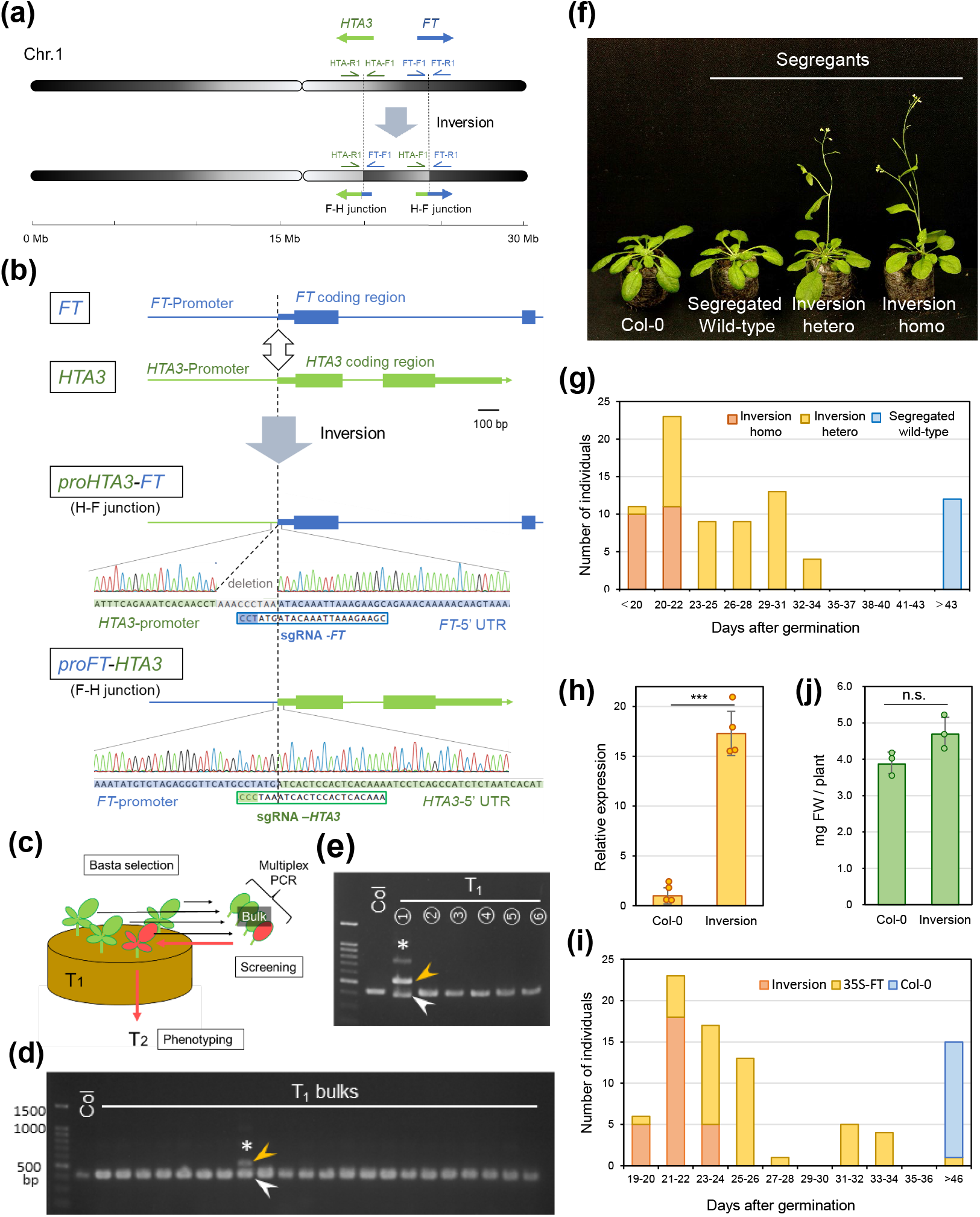
Targeted chromosomal inversion mediated by CRISPR-Cas9 confers an early flowering phenotype in Arabidopsis. **(a)** Diagram of targeted inversion between the *FT* and *HTA3* loci. **(b)** Structures of the fusion sites between *FT* and *HTA3*. Protospacer-adjacent motifs (PAMs) in sgRNAs are shaded in blue or green. Thin and thick boxes indicate untranslated regions (UTRs) and exons, respectively. **(c)** Strategy for screening for the *HTA3-FT* inversion using multiplex PCR-based genotyping. An inversion-positive individual is indicated in red. **(d)** and **(e)**, PCR-based screening for the *HTA3-FT* inversion using genomic DNA of bulked T_1_ individuals **(d)** and individuals from the PCR-positive pool **(e)**. HTA-F1, HTA-R1, and FT-R1 primers (See panel A) were used for multiplex PCR. White and yellow arrowheads indicate the wild-type band and the band expected for the inversion, respectively; * indicates an inversion-positive sample. **(f)** Representative photograph of flowering in Col-0 and inversion segregants grown under short-day conditions at 36 days after germination (DAG). **(g)** Bolting time of transgene-free population segregating for the inversion under short-day conditions. **(h)** *FT* expression levels in Col-0 and inversion homozygotes. Total RNA was extracted from seedlings collected at 9 DAG under short-day conditions. ***, P<0.001 (*n*=4). **(i)** Distribution of bolting time in the transgene-free *HTA3-FT* inversion line and the *CaMV35S:FT* line grown under short-day conditions. **(j)** Fresh weight was measured from the ten 14-day-old plants. n.s., not significant (*n*=3). Statistical analyses were conducted using Student’s *t*-test.

In this strategy, the *HTA3* promoter should be joined to the *FT* coding region (H-F junction), and the *FT* promoter to the *HTA3* coding region (F-H junction). We employed multiplex genotyping PCR that detects both the H-F fusion structure and the F-F wild-type structure (Fig. 1a, c, d). PCR genotyping of bulked DNA samples from 170 independent T_1_ transgenic lines identified the H-F fusion in one bulked DNA sample containing six individuals (Fig. 1d). One of the six individuals showed a positive signal for the H-F fusion (Fig. 1e). We collected T_2_ seeds from this T_1_ plant, genotyped 14 T_2_ plants for the H-F fusion, and examined their flowering time when grown under short-day conditions (Figure S1a). All individuals with at least one copy of the H-F fusion flowered early, whereas individuals that lacked the H-F fusion flowered normally. Thus, the H-F fusion was inherited by the progeny and is associated with early flowering. Sequencing of the PCR fragment derived from the H-F fusion confirmed that the *HAT3* promoter was fused with the *FT* coding region, together with a 9-bp deletion (Figure 1b). For the F-H fusion, sequencing of the corresponding PCR amplicon detected a precise fusion between the *FT* promoter and *HTA3* coding region at the intended break point positions. These findings indicated that an inversion of the 3.6-Mb genomic fragment between *FT* and *HTA3* occurred in this individual.

We generated a CRISPR-Cas9 transgene-free population segregating for the inversion by crossing a plant heterozygous for the inversion with Col-0. We observed a perfect one-to-one correspondence between the presence of an inversion and early flowering, although the flowering phenotype behaved as a semidominant trait (Figure 1f, 1g). The Col-0 plants transformed with a genomic fragment encompassing the inversion exhibited an early flowering phenotype, confirming that the inversion led to earlier flowering (Figure S1b). *FT* is not induced in the wild-type under short-day conditions, but was expressed at high levels in plants homozygous for the inversion under these conditions (Figure 1h). The misexpression of *FT* may therefore be responsible for the early-flowering phenotype of the plants harboring the inversion. RT-PCR indicated that the *FT* transcripts produced in the inversion line comprise their entire respective coding sequences (Figure S2).

When grown under short-day conditions, all transgene-free F_6_ plants homozygous for the inversion flowered as early as transformants overexpressing *FT* from the cauliflower mosaic virus (CaMV) 35S promoter (T_5_ individuals homozygous for a single-locus transgene) (Figure 1i). While the inversion line exhibited a stably inherited early-flowering phenotype, the flowering time of *CaMV35S:FT* plants was more variable with some flowering as late as Col-0, suggesting possible silencing of the transgene. The growth and fertility of the transgene-free line with the inversion were comparable to those of the wild-type (Figure 1j, Figure S3). These observations suggest that the genome-editing inversion system presented here can introduce gain-of-function mutations that confer stable agriculturally important traits, without affecting other traits.

In this study, we successfully modified the expression of a target gene using SDN-1 genome editing in Arabidopsis, a feat that has previously been achieved primarily through creation of genetically modified organisms. This system potentially has interesting applications. For instance, a herbicide-resistant rice strain was developed through overexpression of a target of herbicide by CRISPR-Cas9-mediated inversion (Lu et al., 2021). Beyond overexpression, modifying tissue specificity or enabling the inducible expression of a gene of interest could be achieved by selecting an appropriate promoter. Development of additional technologies, including induction of translocations (Beying et al., 2020), would expand the applicability of this system by enhancing the availability of promoters. Furthermore, the recently reported Bridge RNA system, which achieved programmed recombination in *Escherichia coli*, could be incorporated into this system for the innovative crop improvement (Durrant et al., 2024).

