## Supplemental Figs for "Promoter replacement by genome editing creates gain-of-function traits in Arabidopsis"

a

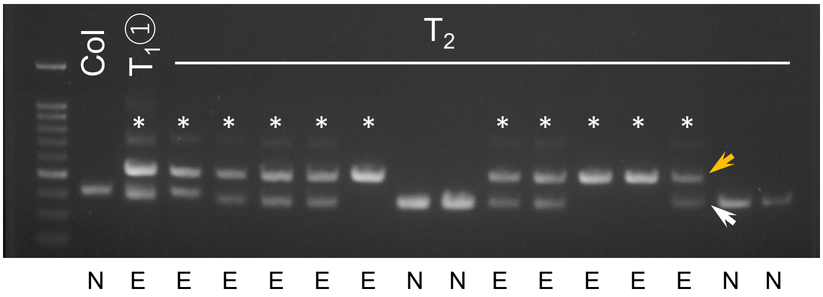


b

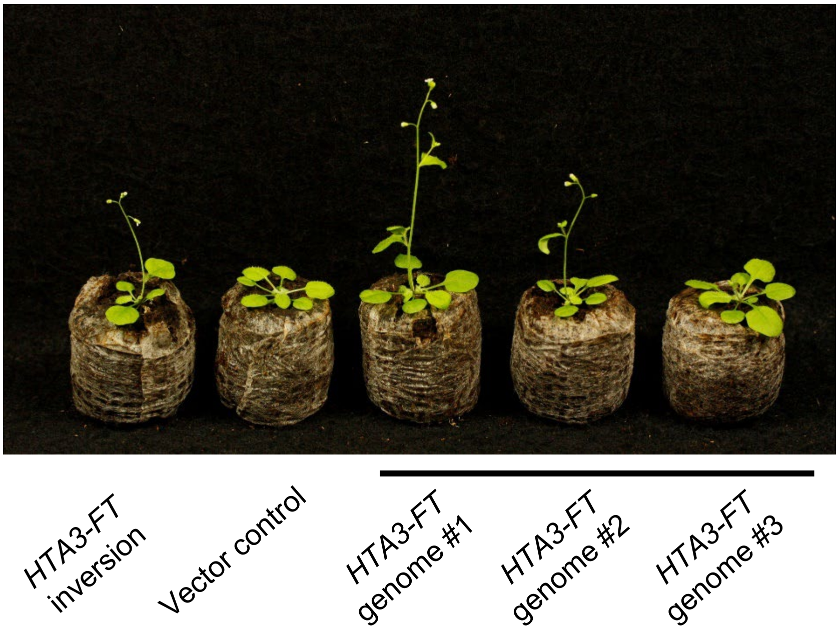


**Figure S1. Early flowering in plants harboring a genomic fragment containing an *HTA3-FT* fusion.** (**a**). Multiplex genotyping PCR; White and yellow arrowheads indicate the wild-type band and the band expected for the inversion, respectively; * indicates an inversion-positive sample. Flowering time was recorded for the $T_2$ individuals derived from the inversion-positive $T_1$ individual under short-day conditions. E, early flowering; N, normally flowering. (**b**) Representative photograph showing the flowering phenotype of 28-day-old $T_1$ transgenic plants harboring an *HTA3-FT* genomic fragment grown under short-day conditions. For comparison, one plant from the genome-edited *HTA3-FT* inversion line is shown. The vector control is a transformant harboring *CaMV35S:Venus*.

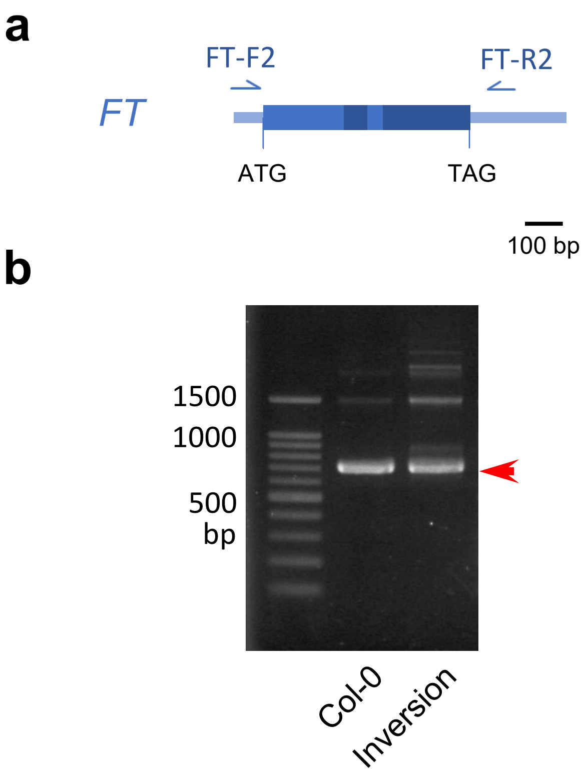


**Figure S2. *FT* expression in the *HTA3-FT* inversion plants. a**, Diagram showing the *FT* mRNA and the positions of primers used for RT-PCR. **b**, RT-PCR products for the *FT* transcript generated using primers designed to anneal to the 5′ UTR and 3′ UTR of *FT*. The arrowhead corresponds to the *FT* transcript encompassing the complete coding sequence, which was verified through Sanger sequencing. Total RNA was extracted from 16-day-old Col-0 seedlings grown under long-day conditions and 9-day-old seedlings of the inversion line grown under short-day conditions.

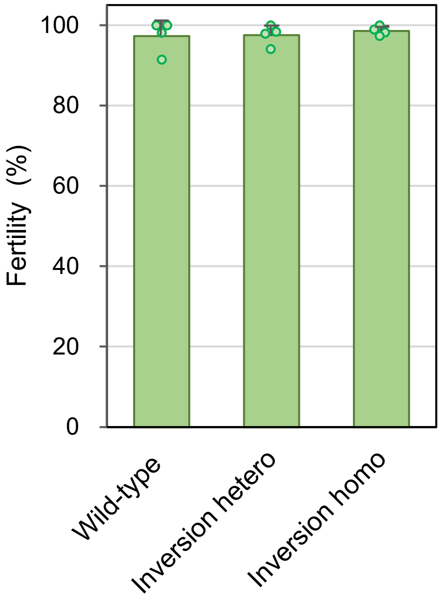


**Figure S3. Fertility analysis of plants homozygous or heterozygous for the *HTA3-FT* inversion.** Fertility was measured as the percentage of placenta that develop seeds in developing siliques. ANOVA revealed no significant differences among the three genotypes (*n*=4).
