## Supplemental Methods for "Promoter replacement by genome editing creates gain-of-function traits in Arabidopsis"

**Methods S1**

**Plant materials, growth conditions, and transformation**

The Arabidopsis accession Columbia-0 (Col-0) was used as the wild-type for transformation. After 4 days of incubation at 4°C in darkness, seeds were sown in Jiffy-7 peat pellets and grown under short-day (10-h light/14-h dark photoperiod) or long-day (16-h light/8-h dark photoperiod) conditions at 22°C and a light intensity of ~100 µmol photons $m^{-2}$ $s^{-1}$. Plants were transformed using the floral-dip method with *Agrobacterium tumefaciens* strain EHA-105. Basta-resistant transgenic individuals were selected following germination and growth on Jiffy-7 peat pellets irrigated with a 50 mg/ml Basta solution.

To obtain a transgene-free population segregating for the inversion, a $T_1$ individual was crossed with Col-0; an $F_1$ individual that did not carry the CRISPR-Cas9 cassette was obtained from the $F_1$ progeny. The $F_2$ seeds collected from this $F_1$ individual were used in the experiment represented in Fig. 2a. $F_3$ seeds from one of the $F_2$ plants homozygous for the inversion were used to establish the *HTA3-FT* inversion line.

Seeds from individual *CaMV35S:FT* $T_1$ transgenic plants were collected. $T_1$ plants with a single-locus T-DNA insertion were identified based on a segregation pattern of 3:1 for Basta resistance in their $T_2$ progeny. $T_2$ lines whose $T_3$ progeny were all resistant to Basta were analyzed as homozygous lines for the transgene.

**PCR screening for the inversion**

Multiplex genotyping PCR was performed using KOD FX Neo DNA polymerase (TOYOBO) with the following three primers: HTA-F1 (5′-AGCTAATTTCGGTTCAACATTGGATG-3′), HTA-R1 (5′-AAGATCGAGAGACGGACTTTGTAG-3′), and FT-R1 (5′-CTCTATAAACTTGGCGGTACCCTAC-3′). A 550-bp amplicon indicated the presence of an inversion (*HTA3-FT*), while a 420-bp amplicon indicated the wild-type configuration (*HTA3-HTA3*). Sequences at the two inverted junctions (*HTA3-FT* and *FT-HTA3*) were determined by Sanger sequencing of the PCR amplicons generated with primers HTA-F1 and FT-R1, and FT-F1 (5′-CTCTATAAACTTGGCGGTACCCTAC-3′) and HTA-R2.

**RNA extraction and quantification**

Seedlings were sampled at zeitgeber 9 (ZT9, with lights on being ZT0) from 9 DAG seedlings grown under short-day conditions and at ZT15 from 16 DAG seedlings grown under long-day conditions. Total RNA was extracted using TRI reagent (Molecular Research Center) according to the manufacturer's instructions. First-strand cDNA was synthesized using a ReverTra Ace qPCR RT Master Mix with gDNA Remover (TOYOBO). Quantitative PCR (qPCR) analysis was performed using the ΔΔ Ct method and a KAPA SYBR FAST qPCR Kit (KAPA Biosystems) on a Rotor-Gene Q 2PLEX thermocycler (Qiagen) with *ACT8* as reference. qPCR primers used were *ACT8* (5′-TGTTGCCATTCAAGCTGTTC-3′ and 5′-TGGAAGTGAGAAACCCTCGT-3′), *FT* (5′-CCCTGCTACAACTGGAACAAC-3′ and 5′-CACCCTGGTGCATACACTG-3′), and *HTA3* (5′-TTAAAGCCGGTAAATACGCCGAAC-3′ and 5′-TTCCAGCTAGCTCCAATACCTCAG-3). Primers used to amplify *FT* transcript in the inversion line were FT-F2 (5'-ACAAATTAAAGAAGCAGAAACAAAAACAAG-3') and FT-R2 (5'-GGCATCATCACCGTTCGTTACTCGTATC-3').

**Fertility and biomass**

To assess fertility, siliques were harvested from Arabidopsis plants about 4 weeks after bolting and the numbers of developing seeds and placentas were counted. To evaluate biomass, the weight of whole shoots of 14-day-old seedlings grown under short-day conditions was measured.

**Author Contribution**

M.K. and T.N. conceived the study and designed the experiments. T.N., and Y.N. performed the experiments. M.N. and M.K. analysed the data.
